# Improved split prime editors enable efficient in vivo genome editing

**DOI:** 10.1101/2024.11.05.622013

**Authors:** Rongwei Wei, Zhenxing Yu, Lihong Ding, Zhike Lu, Keyi Yao, Heng Zhang, Binglin Huang, Miao He, Lijia Ma

## Abstract

Prime editor (PE) is a precise genome-editing tool that enables all 12 possible base-to-base conversions, as well as insertions and deletions, that does not require DSBs or donor DNA. The efficient delivery of prime editors in vivo is critical for realizing the full potential of prime editing in disease modeling and therapeutic correction. Although PE has been divided into two halves and delivered using dual adeno-associated viruses (AAVs), editing efficiency at different gene loci varies among split sites, and efficient split sites within Cas9 nickase are limited. In this study, by screening multiple split sites, we demonstrated that 1115 (Asn) is an efficient split site when delivering PE by dual-AAV. Besides, we utilized a feature reported by others recently that RNase could be detached from the Cas9n and designed split sites in the first half of Cas9n. We found that split-PE-367 enabled high prime editing efficiency with Rma intein. To test the editing efficiency in vivo, dual-AAV split-ePE3-367 was packaged in AAV9 and delivered by tail vein injection in mice, achieving 24.4% precise genome editing 4 weeks after injection. Our findings establish an alternative split-PE architecture that could achieve robust gene editing efficiency, facilitating the potential utility both in model organisms and as a therapeutic modality.

## INTRODUCTION

Prime Editors (PEs) are precise genome editing tools that enable all types of base conversions, small deletions, and insertions(Anzalone et al. 2019). PEs minimally require a Cas9 nickase (Cas9n) and a reverse transcriptase (RT) paired with a prime-editing guide RNA (pegRNA). The pegRNA carries a single guide RNA (sgRNA) and a 3’ extension that consists of an RT template encoding the desired modifications and a prime binding site (PBS). Upon binding to the target site, the Cas9n nicks the non-target strand, and the RT uses the RT template to generate complementary DNA for precise repair of the nicked site. PEs incorporate desired modifications without the requirement of double-strand breaks (DSBs), thus reducing nuclease-associated unwanted editing outcomes such as large deletions(Kosicki et al. 2018), inversions(Mani and Chinnaiyan 2010), translocations(Kosicki et al. 2018), and the initiation of the p53 DNA damage response(Haapaniemi et al. 2018; Ihry et al. 2018; Enache et al. 2020). Moreover, PEs have been found to induce very few detected off-target genomic changes in vitro and in vivo(Liu et al. 2020; Jin et al. 2021; Lin et al. 2021; Park et al. 2021). Therefore, the ability to precisely install or correct pathogenic mutations independent of their composition makes prime editing a promising tool to perform somatic genome editing in model organisms to study disease processes or to utilize for therapeutic applications.

The editing efficiency of the initial PE2 system is modest. Therefore, an additional sgRNA is used to nick the non-edited strand (PE3), improving the editing efficiency by 1.5- to 4.2 folds over PE2(Anzalone et al. 2019). Besides, proteins that could improve DNA repair and Chromatin accessibility were also applied(Chen et al. 2021; Park et al. 2021; Ferreira da Silva et al. 2022). These strategies successfully improve the editing efficiency, but the size of PE is larger as well, making it difficult for safe and efficient delivery of PE in vivo(van Haasteren et al. 2020). Owing to its relatively low immunogenicity profile and efficient cellular undertaking, adeno-associated virus (AAV) has been the most commonly used vector in the therapeutic application(Varnavski et al. 2005; Verdera et al. 2020). Unfortunately, PE (>6.3 kb) exceeds the packaging capacity of AAV (∼4.7 kb). To address this issue, PE is divided into two smaller parts that are separately fused to one-half of the split-intein and co-delivered by a dual-AAV system(Liu et al. 2021a; Zheng et al. 2022; Zhi et al. 2022; She et al. 2023). Then, the two halves could be reassembled into full-length PE by a trans-splicing intein at the post-translational level. Previous studies suggest that different split sites affect structural stability and, therefore, the constituting activity, resulting in different genome-editing efficiency(Li 2015). Efforts have been taken to screen split sites for the improvement of gene editing efficiency of split-intein PE systems in vivo, and they have achieved maximum efficiencies corresponding to 1.7% - 13.5% editing of bulk liver or retina(Liu et al. 2021a; Böck et al. 2022; Gao et al. 2022; Jang et al. 2022; Zheng et al. 2022; Zhi et al. 2022; She et al. 2023). Recently, by increasing PE expression, prime editing guide RNA stability, and modulation of DNA repair, David R. Liu developed optimized split-PE-1024 vectors (v3em PE3-AAV) and achieved up to 46% editing efficiency in mouse liver using dual-AAV(Davis et al. 2024). These studies indicate that dual-AAV split-PE is a feasible tool for animal research and therapeutic applications. However, due to the packaging limit of AAV, Cas9n can be only split in the second half to ensure that the C-terminal fragment of PE (C-terminal of Cas9n and RT) could be packaged with a promoter and a poly(A) signal into a single AAV vector. Recent studies show that separated Cas9n and MMLV-RT protein function as efficiently as intact PE2(Liu et al. 2022; Grünewald et al. 2023), making it possible to screen split sites at the second half of Cas9n and construct more efficient split-PEs.

In this study, we tested various split sites in vitro and found that split-PE-1115 exhibited high editing efficiency. Besides, by using untethered RT, we screened split sites located at the second half of Cas9n and identified an efficient split-PE (split-PE-367) with robust editing efficiency. We further investigated the efficiency of the two split PEs (split-PE-1115 and split-PE-367) at multiple genomic loci and observed the successful installation of short insertions and point mutations at these loci. Finally, we packaged split-ePE3-367 into AAV9 to target *Pcsk9*, a gene involved in cholesterol homeostasis. We showed that the delivery of split-ePE3-367 by dual-AAV9 realized the installation of the mouse homolog of *PCSK9* Q152H with up to 24.4% efficiency in a precise way, demonstrating that split-ePE3-367 is an alternative split PE that displays high editing efficiency in vivo.

## RESULTS

### Screening of split sites for efficient PE delivery

PE2 contains an engineered MMLV-RT (2031 bp) fused to the C-terminal of SpCas9 (H840A) nickase (SpCas9n). To ensure that each split segment of PE would not exceed the packaging limit of an AAV vector, we first screened split sites within the range from amino acids 855 (Val) to 1155 (Lys) of SpCas9. Meanwhile, the C-intein should be placed in front of a nucleophilic amino acid (cysteine, threonine, or serine) to achieve high splicing efficiency(Li et al. 2008).

According to these two criteria, we selected four split sites to deliver PE2 by dual-AAV vectors (Fig. 1a). One of the most commonly used inteins in split-PE, the Rma intein from *Rhodothermus marinus* DnaB, was employed in this study. At the post-transcriptional level, the two flanking peptides can be ligated through intein-mediated protein trans-splicing (Fig.1b). We assessed the prime editing efficiency of different split-PEs at the *EMX1* gene locus (EMX1_5_G > T) in HEK293T cells. 72 h post-transfection, cells were harvested for genomic DNA extraction, and the editing efficiency was assessed by next-generation sequencing (NGS). The results showed that splitting PE2 after amino acid 1115 (referred as split-PE2-1115) mediated the most efficient on-target editing among the four split sites (Fig.1c). Split-PE2-1115 improved prime editing efficiency by 2.5 folds compared to split-PE2-1024, an efficient split PE that has been verified(Zhi et al. 2022). Full-length PE2 expression by split-PE2-1115 and split-PE2-1024 was confirmed by western blotting, and we found higher expression of PE by split-PE2-1115, which is consistent with the results of NGS (Fig. S1).

**Figure 1.**
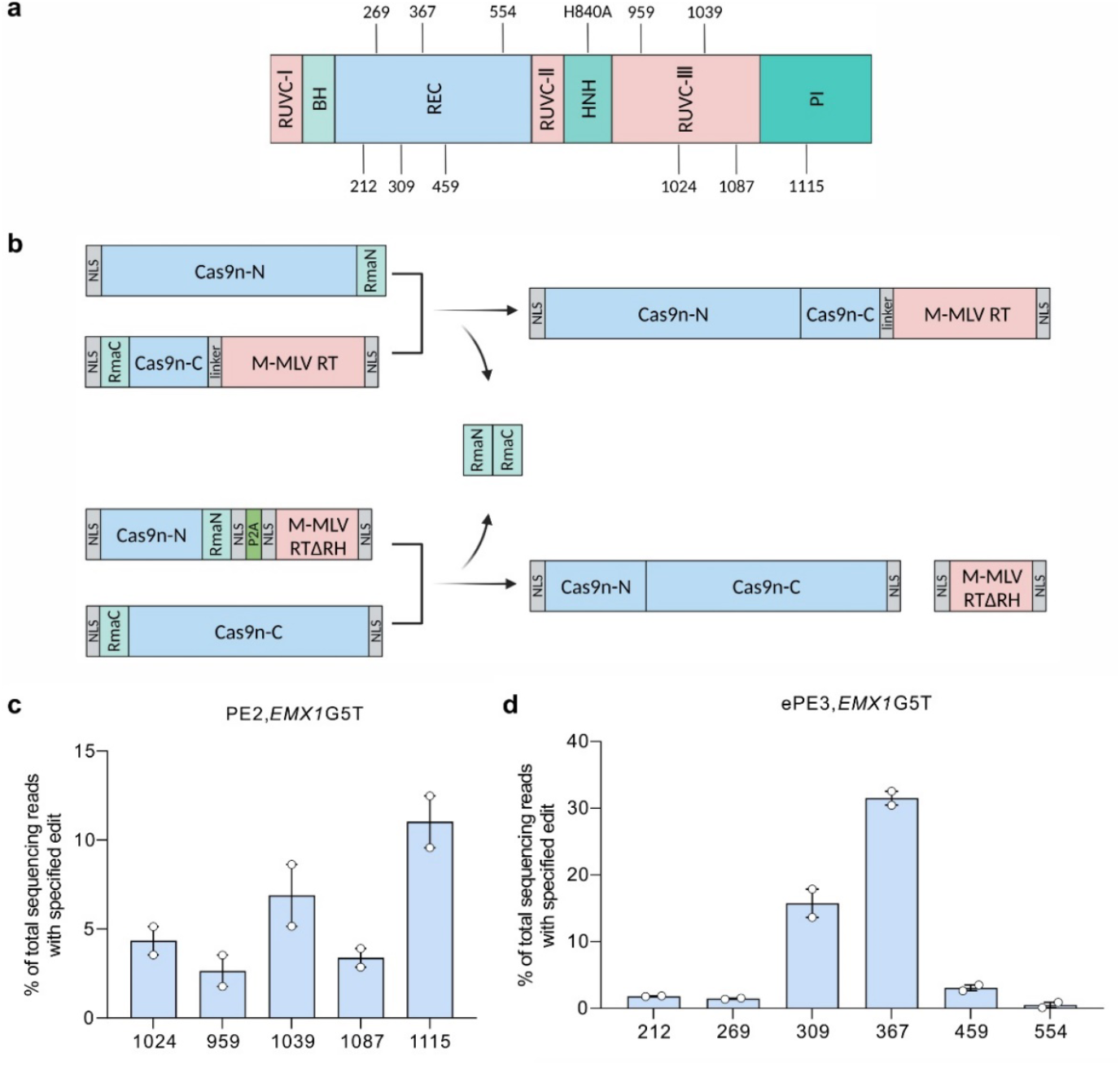
Development of efficient split-PE in HEK293T cells. The multisite split-PEs induce a single-base conversion at the endogenous loci. **a**, Schematic of SpCas9n in the PE fusion protein splits at 11 different sites (212-213, 269-270, 309-310, 367-368, 459-460, 554-555, 959-960, 1024-1025, 1039-1040, 1087-1088, 1115-1116). RuvC, endonuclease domain; BH, bridge helix; REC, recognition domain; HNH, His-Asn-His endonuclease domain; PI, protospacer-adjacent motif (PAM)-interacting domain. **b**, Schematic of Rma intein-mediated PE or SpCas9n reconstitution through protein trans-splicing. NLS, nuclear localization sequence. **c**, Screening of split-PE2s with different split sites at the second half of SpCas9n. The frequencies of total sequencing reads with specified editing introduced by different split-PEs were quantified by NGS and analyzed by CRISSPResso2 (https://github.com/pinellolab/CRISPResso2). HEK293T cells were transfected with split-PE2 plasmids at equal molar concentrations. Data are shown as mean ± SEM, n = 2. **d**, Screening of split-ePE3s with different split sites at the first half of SpCas9n. P2A peptide was used to co-express an untethered MMLV-RTΔRH protein. The frequencies of total sequencing reads with specified editing introduced by different split-PEs were quantified by NGS and analyzed by CRISSPResso2 (https://github.com/pinellolab/CRISPResso2). HEK293T cells were transfected with split-ePE3 plasmids at equal molar concentrations. Data are shown as mean ± SEM, n = 2.

Next, an additional nicking sgRNA and an engineered pegRNA (epegRNA) with a trimmed evopreQ1 motif to the 3′ terminus were used to improve PE editing efficiency(Nelson et al. 2022) (ePE3). Besides, a truncated MMLV RT (MMLV-RTΔRH), which is encoded by 1,488 bp and is 26.7% smaller than the parental MMLV RT(Grünewald et al. 2023), was employed to minimize the size of PE. What’s more, we introduced tRNA(Yuan and Gao 2022) or hammerhead ribozyme (HH)(Tiyaboonchai et al. 2022) to further reduce the size of PE. We assembled the nicking sgRNA downstream of the epegRNA followed by a tRNA sequence or a HH sequence, which can be processed and subsequently release individual nicking sgRNA and epegRNA. Three mature tRNAs (RNase fully processed tRNA sequence) were selected (hGly GCC, hPro AGG and hGln CTG)(Shiraki and Kawakami 2018). The results showed that the efficiency of both the tRNA and the HH architectures decreased compared to that of U6, demonstrating insufficient expression of epegRNA or nicking RNA (Fig. S2).

Previous studies revealed that the RT activity is likely provided by a second PE molecule that is not bound to the target DNA site and SpCas9n with an untethered RT function similarly to intact PE2 protein(Liu et al. 2022; Grünewald et al. 2023). We thus replaced the XTEN linker between the C-terminal of SpCas9n and RT with P2A, resulting a dissociative RT (Fig. S3a). We probed the editing efficiency of ePE3 split after amino acid 1115 with an untethered MMLV-RTΔRH (split-ePE3-1115) and found that ePE3 with untethered MMLV-RTΔRH was just as efficient as that with tethered MMLV-RTΔRH (Fig. S3b). The result implies that the MMLV RT and the N-terminal of SpCas9n can be encoded in one AAV vector. This broadens the split window and allows SpCas9n to be split up to amino acid 203.

To identify more optimal split sites, we rearranged the dual-AAV vectors, one encoding the N-terminal of SpCas9n and MMLV-RTΔRH(Zheng et al. 2022) and the other encoding the C-terminal of SpCas9n (ePE3)(Anzalone et al. 2019; Nelson et al. 2022) (Fig. 1b).The epegRNA and nicking sgRNA cassettes were assembled together or separately to one of the AAV vectors based on the cargo size of AAV. To further increase the editing efficiency and provide more flexibility to accommodate the essential components of PE, we adapted various strategies. For example, a shorter poly A signal, SV40 poly A signal, was used to substitute the bGH poly A signal when the overall packaging size exceeds the capacity of AAV. Moreover, a previous study indicated that expressing two gRNAs from two identical U6 promoters is prone to recombination and consequent loss of the proximal gRNA(Vidigal and Ventura 2015). Thus, we incorporated a tRNA to express epegRNA and nicking sgRNA from the same expression cassettes when they were encoded in the same vector (Fig. S4). Together, we tested six different split sites and found that split-ePE3-367 showed the highest precise editing efficiency among the six split-ePE3 variants (Fig. 1a), realizing 31.5% G > T editing at *EMX1* locus (Fig. 1d). Furthermore, we tried to find more effective split sites around the split-367. The data showed that split-ePE3-354 exhibited similar editing efficiency to that of split-ePE3-367 while split-ePE3-385 showed decreased editing efficiency (Fig. S5).

Next, we tried to optimize split-PE3-367. Since the orientation and position of the gRNA relative to the Cas9 might affect both Cas9 and gRNA expression, further influence the gene editing efficiency(Ran et al. 2015; Fry et al. 2020; Levy et al. 2020). We generated different AAV vectors by changing the orientation and positions of epegRNA and nicking sgRNA (Fig. S6a). Besides, we also located the MMLV-RTΔRH upstream of the N-terminal of Cas9 to construct a new split-PE architecture. Compared to the initial split-ePE3-367, all the architectures showed comparable editing efficiency except split-ePE3-367_V3, which resulted in a slight increase of prime editing efficiency (Fig. S6b).

### Efficient genome editing by split-PEs at multiple endogenous sites in human cells

Then we assessed the editing efficiency of split-ePE3-1115 and split-ePE3-367 at multiple endogenous sites, and the published split sites applicable for verifying in vivo applications (split-ePE3-1105, split-ePE3-1024, and split-ePE3-1024CFN) were included as controls(She et al. 2023; Davis et al. 2024). First, we selected five commonly targeted endogenous genes and tested the editing efficiency. We transfected plasmids encoding two halves of ePE3 along with epegRNA and nicking sgRNA into HEK293T cells and quantified the editing efficiencies 72 h post-transfection by NGS. Both the split-ePE3-367 and split-ePE3-1115 exhibited similar or higher editing efficiency than split-ePE3-1105, split-ePE3-1024 and split-ePE3-1024CFN (Fig. 2a). Then we used split-ePE3 to install therapeutically relevant mutations associated with cardiovascular disease, type 2 diabetes, Alzheimer’s disease, Rett syndrome and transthyretin amyloidosis. We observed that both split-ePE3-1115 and split-ePE3-367 achieved robust prime editing efficiency, comparable to that of the previous split-PEs (Fig. 2b). Besides, across the eight endogenous sites we tested for single-base substitutions, split-ePE3-367 demonstrated comparable or higher editing efficiencies compared to the split-ePE3-1024 and split-ePE3-1024CFN (Fig. 2a, b). At three of the four endogenous sites we tested for small insertions, the insertion efficiencies of split-ePE3-1115 were superior to others (Fig. 2a, b). Collectively, these data demonstrated that PEs split at both 367 and 1115 could install single-nucleotide transversions and small insertions at endogenous sites with high efficiency, indicating that the 367 and 1115 were two promising split sites for dual-AAV delivery of PE.

**Figure 2.**
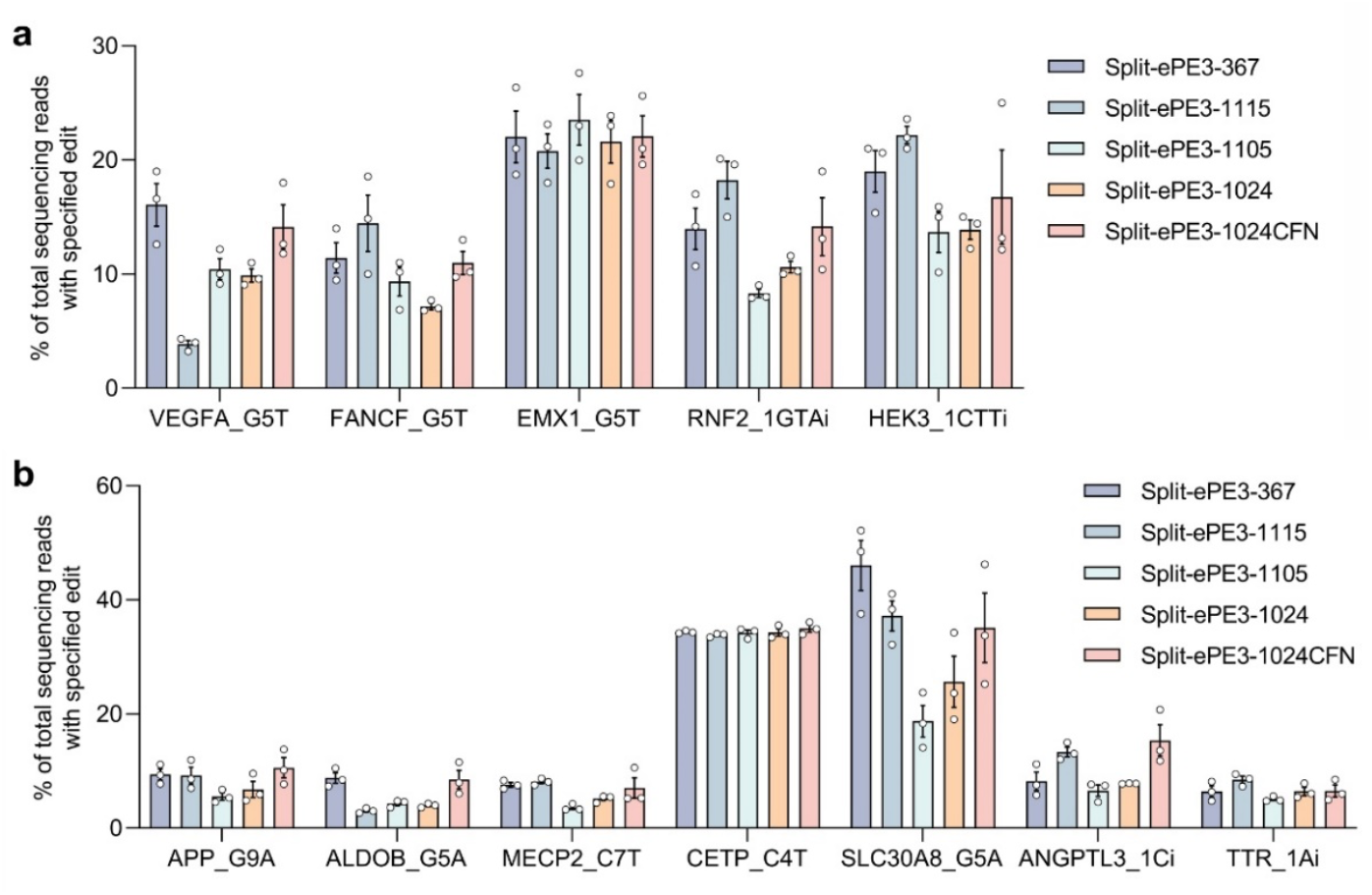
Comparison of the editing efficiency of five split-PEs at multiple endogenous sites. **a**, Precise editing efficiency of the five split-PEs at five commonly used endogenous sites. **b**, Efficiency of the five split-PEs installing therapeutically relevant mutations at seven endogenous sites. Split-ePE3-1105, split-ePE3-1024, and split-ePE3-1024CFN were three published split-PEs with high gene editing efficiency. The frequencies of total sequencing reads with specified editing introduced by different split-PEs were quantified by NGS and analyzed by CRISSPResso2 (https://github.com/pinellolab/CRISPResso2). HEK293T cells were transfected with split-ePE3 plasmids at equal molar concentrations. Data are shown as mean ± SEM, n = 3.

### Delivery of split-ePE3-367 using AAV for in vivo prime editing

We have demonstrated that split-ePE3-367 mediated high-efficiency installation of single-base substitutions in human cells by plasmid transfection. We next investigated whether split-ePE3-367 could realize efficient gene editing in vivo using dual AAVs. Here, we targeted mouse Proprotein convertase subtilisin/kexin type 9 *(Pcsk9)* gene and installed a G-to-C substitution that blocks autocatalytic processing of Pcsk9(Mayne et al. 2011) (Fig. 3a). PCSK9 is a proprotein convertase that involved in the degradation of low-density lipoprotein (LDL) receptor, a key receptor that mediates endocytosis of cholesterol, and has been a therapeutic target for cardiovascular disease(Abifadel et al. 2003; Cohen et al. 2005; Seidah 2021; Libby and Tokgözoğlu 2022). To improve PE expression in mice, we used the Cbh promoter, a ubiquitous and constitutive promoter that is less sensitive to silencing(Gray et al. 2011) in vivo, to replace the CMV promoter (Fig. 3b). Besides, a truncated SV40 poly(A) was applied to minimize the size of transgenes in AAV(Gimmi et al. 1988). We first tested on-target editing efficiency at the endogenous *Pcsk9* locus in Hepa1-6 cells by plasmid transfection and observed 22.7% G-to-C substitution, similar to that of v3em PE3-AAV (25.7%)(Davis et al. 2024) (Fig. 3c).

**Figure 3.**
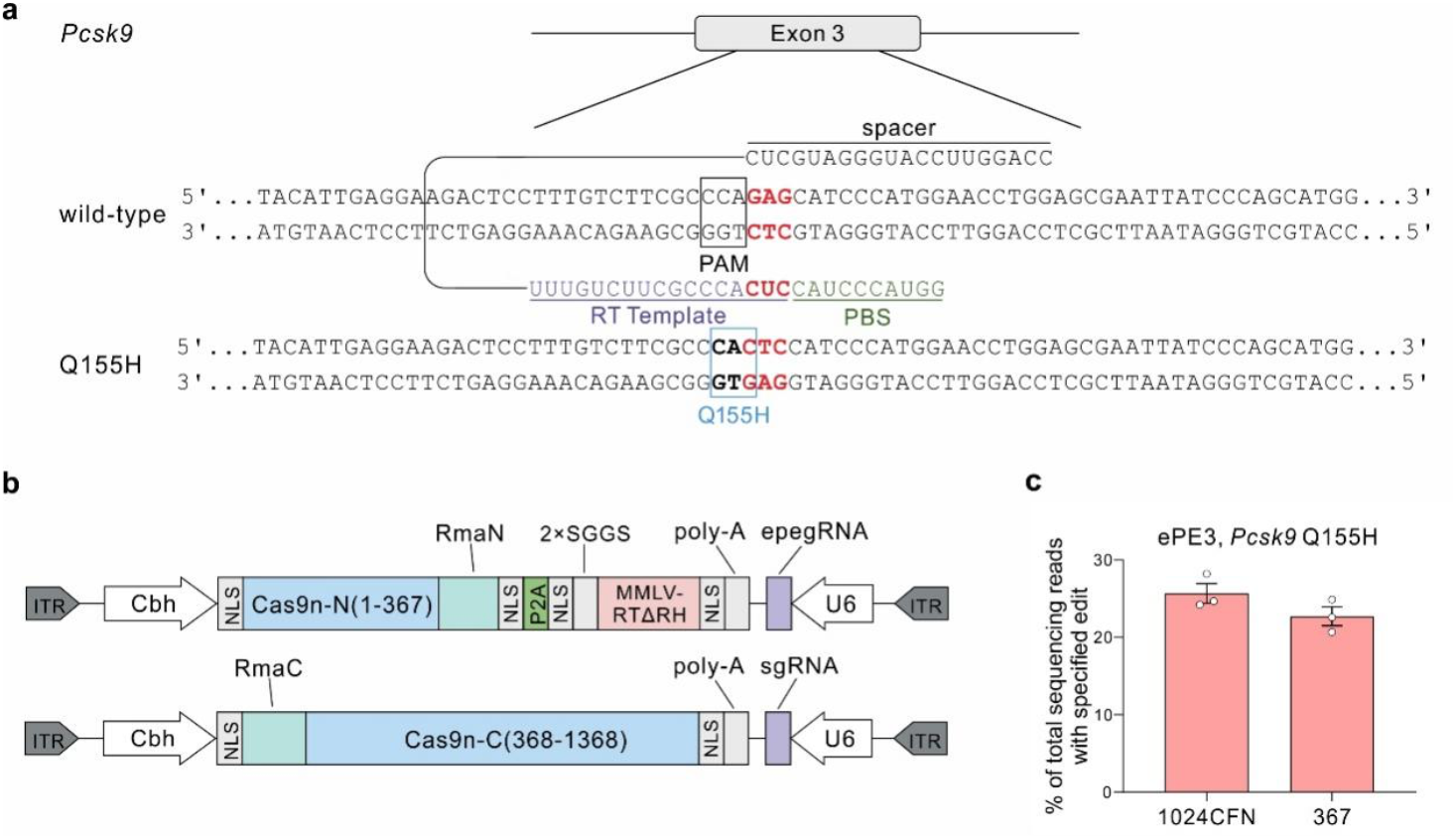
Split-ePE3-367 mediated installation of m*Pcsk9* Q155H in Hepa1-6 cells. **a**, Schematic of installation the mouse homolog of *Psck9* Q152H using Split-ePE3-367. **b**, Schematic of the AAV vector genomes for split-ePE3-367. **c**, Editing efficiencies of split-ePE3-367 and split-ePE3-1024CFN for installation of *Psck9* Q155H in Hepa1-6 cells. The frequencies of total sequencing reads with specified editing were quantified by NGS and analyzed by CRISSPResso2 (https://github.com/pinellolab/CRISPResso2). Hepa1-6 cells were transfected with split-ePE3 plasmids at equal molar concentrations. Data are shown as mean ± SEM, n = 3.

We next characterized the performance of split-ePE3-367 in vivo. The two vectors of split-ePE3-367 were packaged into AAV9 and a total dose of 2×10^12^ vector genomes (vg) (1×10^12^ vg each of the N-terminal half of SpCas9n/MMLV-RT/epegRNA and the C-terminal of SpCas9n/nicking sgRNA AAVs) was injected into 6-week-old mice through the tail vein (Fig. 4a). Three weeks after injection, genomic DNA was isolated from the liver, and the target locus was analyzed by NGS. The results showed that split-ePE3-367 achieved 24.4% precise editing (Fig. 4b). To assess the off-target effects of split PE in vivo, we performed NGS at 10 potential off-target loci that might be targeted by pegRNA. No detected off-target was observed at these loci in mice treated with split-ePE3-367 AAV9 (Fig. S7), indicating high-specificity editing of split PE at this site. Overall, these results demonstrate that split-ePE3-367 can mediate robust and specific therapeutically relevant prime editing in vivo using a dual-AAV system.

**Figure 4.**
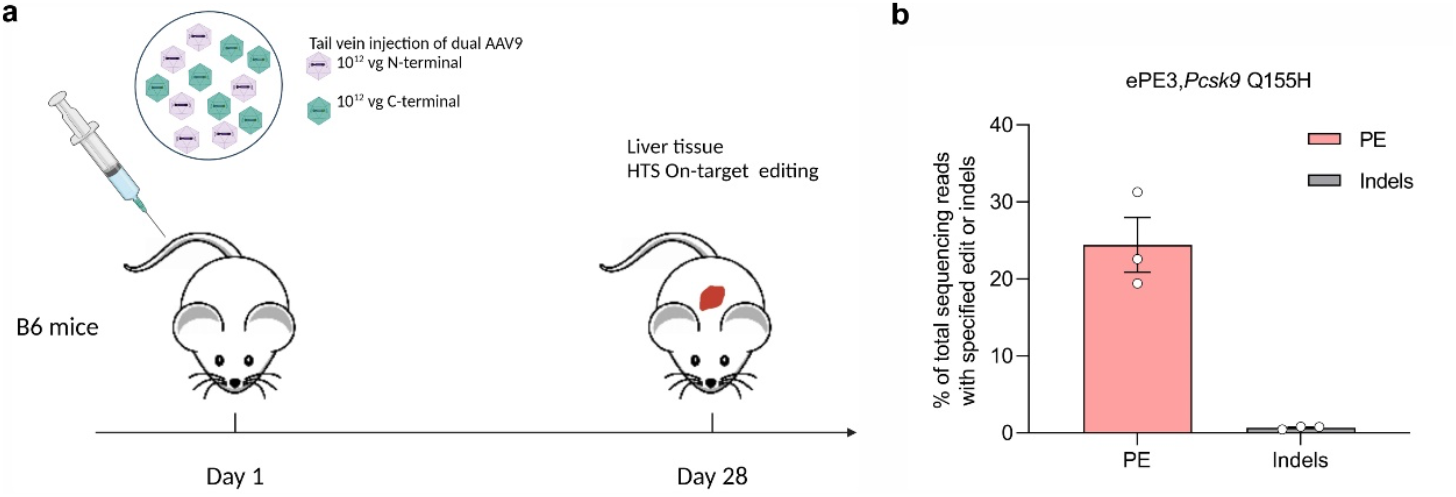
*In vivo* prime editing to install *Pcsk9* Q155H by split-ePE3-367. **a**, Schematic of the tail-vein injection experiments with dual-AAV split-ePE3-367. Six-week-old C57BL/6 mice were injected with a total of 2×10^12^ vg AAV9. After 3 weeks, liver genomic DNA was sequenced. **b**, Editing efficiencies of split-ePE3-367 for installation of *Psck9* Q155H in mouse.

## DISCUSSION

PE is the first precise genome-editing technology and its broad spectrum potentially allows the correction of up to 89% of human genetic diseases(Zhao et al. 2023). Many strategies have been developed by engineering PE protein and pegRNA to improve prime editing efficiency(Chen et al. 2021; Liu et al. 2021a; Liu et al. 2021b; Park et al. 2021; Song et al. 2021; Böck et al. 2022; Gao et al. 2022; Li et al. 2022; Nelson et al. 2022; Velimirovic et al. 2022; Zhang et al. 2022; Zheng et al. 2022). Recently, PE7 has been developed by fusing LA, a small RNA-binding protein, and effectively enhanced prime editing efficiency in vitro (Yan et al. 2024). However, due to the large size of PE, effective delivery of it in vivo using AAV remains a challenge. Dual-AAV vectors have been used for in vivo PE delivery by dividing PE into two halves. Since the choice of split sites affects editing efficiency, we screened four split sites using PE2 and found split-PE2-1115 showed the highest on-target efficiency at *EMX1*. What’s more, we applied an untethered MMLV-RTΔRH, which was encoded together with the N-terminal of Cas9n in one AAV vector, and thus screened more split sites within a range from amino acid 203 to 1368. We found that split-ePE3-367 mediated robust G-to-T conversion at *EMX1*. Another split site near 367 also exhibited high editing efficient. We further verified the editing efficiency of split-PEs (split-ePE3-1115 and split-ePE3-367) at multiple endogenous sites. Finally, we delivered the optimized split-ePE3-367 into mice using AAV9 and achieved 24.4% precise editing at *Pcsk9*.

The editing efficiency of split PE at different gene loci varies among split sites. It’s necessary for us to discover more robust split sites to realize efficient prime editing at specific loci which still be edited with modest efficiency. Previous studies have demonstrated that split-PE2-1115 resulted in decreased on-target editing compared to split-PE2-1105(She et al. 2023). However, we tested the editing efficiency of split-ePE3 separately divided after 1115 and 1104 at various gene loci and found that split-ePE3-1115 performed better at most of the loci (Fig. 2), indicating that 1115 is also a feasible split site when deliver PE using dual AAV. Besides, split-ePE3-367 with untethered MMLV-RTΔRH also exhibited high prime editing efficiency. A previous study showed that PE allows efficient gene editing when delivered by two AAV vectors which separately encode intact SpCas9n and MMLV-RT without using inteins(Grünewald et al. 2023). However, this was achieved by strict limitation in the promoter length due to the large size of SpCas9n. Delivering of PE with a split SpCas9n and untethered RT provides a novel split-PE architecture that enables the utility of longer promoters to drive the PE expression. Meanwhile, it also broadens the range among which more split sites can be found. By screening split sites near amino acid 367, we also found another split site that could realize high primer editing efficiency. In conclusion, we have demonstrated more alternative split sites that can be applied to efficiently deliver PE in vivo by AAV. Furthermore, base editors are another gene editing tool that could catalyze highly efficient base deamination-induced base transitions without producing DSBs(Anzalone et al. 2020; Porto et al. 2020). In base editors, the DNA deaminase enzyme is usually fused to the N-terminal of Cas9 nickase, which could only be split from amino acid 179 to 740 in the SpCas9n domain when delivered using AAV(Villiger et al. 2018; Chen et al. 2020; Levy et al. 2020). Therefore, the efficient split sites close to the N-terminal of Cas9n, such as 367 and 354, provide a potential site for dividing base editors as well.

## Methods

### Cell culture, transfection and genomic DNA extraction

HEK293T cells and Hepa1-6 cells were cultured in high-glucose Dulbecco’s modified Eagle’s medium (DMEM) supplemented with 10% fetal bovine serum (FBS) (Gemini #900-108) and 1% GlutaMAX (Thermo #35050061). Cells were grown at 37°C with 5% CO_2_ and passaged every 2-3 days when cells reached approximately 80% confluency. Cells were transfected at 60%-70% confluency using 3 uL of Lipofectamine 3000 (Thermo Fisher) with 1500 ng each half of the Split-PE. Three days post transfection, cells were collected and then lysed to extract genomic DNA using TIANamp Genomic DNA Kit (TIANGEN).

### Deep sequencing and data analysis

Deep sequencing of genomic DNA was performed as previously described(Anzalone et al. 2019). For the first round of PCR, genomic sites of interest were amplified from 150 ng of genomic DNA using Q5 PCR Master Mix (New England Biolabs #M0544S) with the primers containing Illumina forward and reverse adapters (listed in Table S1). 50 uL of PCR 1 reaction was performed with forward and reverse primers each at 0.25 uM, ∼1 uL of genomic DNA extract, and 25 uL of Q5 PCR Master Mix. To add a unique sequencing index, 50 uL of a given PCR 2 contained 2.5 uL of previous step unpurified PCR product; a unique forward or reverse primer pair, each at 0.25 uM; and 25 uL of Q5 PCR Master Mix. Primers are listed in Table S1. PCR 2 products were purified by DNA AMPure XP beads (Beckman Coulter #A63882) and eluted with 25 uL of water. DNA concentration was quantified by Nanodrop One (Thermo Fisher Scientific). The library was sequenced using an Illumina NovaSeq system. Deep sequencing data were processed using CRISPResso2 (https://github.com/pinellolab/CRISPResso2). Prime editing efficiency was calculated as the percentage of the reads representing correct editing and the total reads.

### Western blotting

Cells were lysed 72 hours after transfection using cold RIPA buffer supplemented with Phenylmethanesulfonyl fluoride at 1 mM. Cell lysates were quantified using the bicinchoninic acid assay (BCA, Takara). For each sample, 25 ug of protein was loaded onto a 12-well 4%– 20% Bis-Tris Protein Gel, run at 180 V for 1 hour, then transferred to the PVDF membrane (Sigma). The primary antibodies anti-beta-actin (Cell Signaling Technology, dilution: 1:1,000) or anti-Cas9 (Cell Signaling Technology, dilution: 1:1,000) were used, followed by incubation with secondary antibodies. Bands were visualized using the iBright750 imaging system (Thermo).

### Production of AAV

HEK293T cells were transfected with the AAV genome, pHelper, and Rep/Cap plasmids using polyethyleneimine transfection (PEI MAX, Polysciences) at approximately 90% confluency. Three days after transfection, cells were harvested by cell scrapers (Corning), collected by centrifuging at 3000 rpm for 30 min, resuspended in 2 mL lysis buffer every five plates (50 mM Tris-HCl, 500 mM NaCl) and stored at -80°C overnight. Media were decanted to a new bottle with a final concentration of 8% PEG8000 (Sigma) and 500 mM NaCl, incubated at 4°C overnight and then collected by centrifuging at 3400g for 15 min, resuspended in an appropriate amount of cold DPBS, and finally stored at 4°C temporarily. Cell lysates were mixed with previous step product after freeze-thaw 3 times and added Benzonase to the cell lysates at a final concentration of 50 U/ml. After incubating at 37 °C in a water bath for 30 min, media was collected by centrifuging at 5000g for 30 min. During the incubation, an iodixanol gradient was prepared. Then AAV was purified by ultracentrifugation. The AAV solution buffer was exchanged for cold DPBS with 0.001% PF-68 using 100 kDa MWCO columns (Sigma) and concentrated. The titer of the AAV was quantified via qPCR.

### Animals

All animal experiments were conducted under the protocol approved by the Animal Care and Ethical Committee of the Westlake University. C57BL/6 and ICR mice were purchased and housed in the Laboratory Animal Resource Center (LARC) at the Westlake University. The LARC is a certified pathogen-free and environmental-control facility (21±2°C, 55±15% humidity and 12:12-h light : dark cycle).

AAV was diluted into pre-chilled PBS to a concentration of 2 × 10^13^/ml. Subsequently, 125 μl of CBH608 and 125 μl of CBH621 were mixed thoroughly. Using a sterile 0.6 × 25 mm TWLB syringe, 200 μl of the mixture containing 1 × 10^12^ of each virus, were injected into the tail vein of six-week-old male C57BL/6 mice. This experiment was repeated three times independently. After three weeks, the mice were euthanized using carbon dioxide, and their livers were dissected. The livers were washed thoroughly with pre-chilled PBS, minced with surgical scissors, and placed in 2 mL sterile Eppendorf tubes. Two 4 mm and two 2 mm stainless steel beads were added to each tube. The tissues were homogenized using the Qiagen Tissuelyser II at a power setting of 30 for 3 minutes, followed by reversing the adapter orientation and homogenizing again at the same power setting for an additional 3 minutes. Portions of the homogenized tissue were used for DNA extraction. Genomic DNA (gDNA) from the organs was extracted using the TIANamp Genomic DNA Kit (TIANGEN #DP304-03) according to the manufacturer’s instructions.

### Off-target analysis

Potential off-target sites were amplified by PCR using the primer sequences (Table S1), then sequenced on an Illumina NovaSeq system. Deep sequencing data analysis was processed using CRISPResso2 and untreated cells were used as negative controls to exclude PCR amplification and sequencing errors. Any aligned reads with a nucleotide sequence within prime editing target window that matches the nucleotide encoded by the pegRNA reverse transcription were noted as off-target reads. Off-target editing efficiencies, which are greater than 0.02%, were quantified as a percentage of (number of off-target reads) / (number of total reads).

### Statistical analysis

Statistical analyses were performed using GraphPad Prism. Data were represented as biological replicates and mean ± SEM was indicated in the corresponding figures. Likewise, the respective figure legends described sample sizes and the statistical tests used in detail.

