## Supplemental figures for "Improved split prime editors enable efficient in vivo genome editing"

Supplementary Figure Legends S1-S7
Supplementary Tables S1-S2

**SUPPLEMENTARY FIGURES**

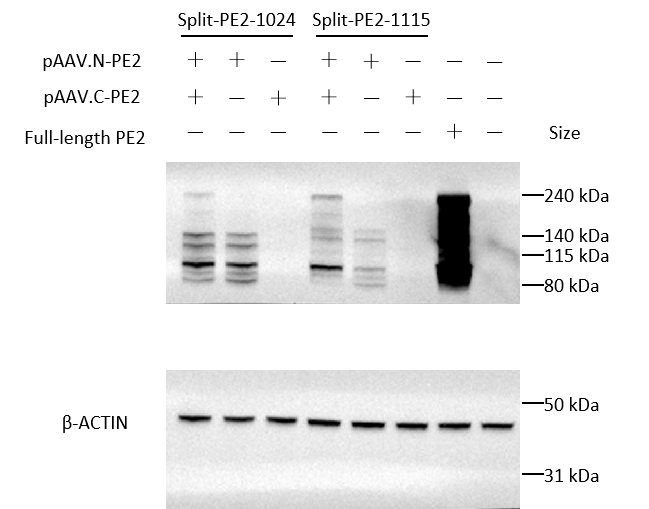

Figure S1. Western blot analysis to detect full length PE expression in HEK293T cells co-transfected with the dual-AAV vectors encoding split-PE2-1115 or split-PE2-1024. Beta-actin was used as a loading control.

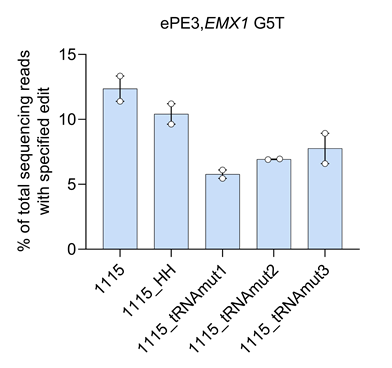

Figure S2. Optimization of split-ePE3-1115 by replacing U6 with hammerhead ribozyme (HH) or mature tRNAs. The frequencies of total sequencing reads with specified editing were quantified by NGS and analyzed by CRISSPResso2 (<https://github.com/pinellolab/CRISPResso2>). HEK293T cells were transfected with split-ePE3 plasmids at equal molar concentrations. Data are shown as mean ± SEM, n = 2.

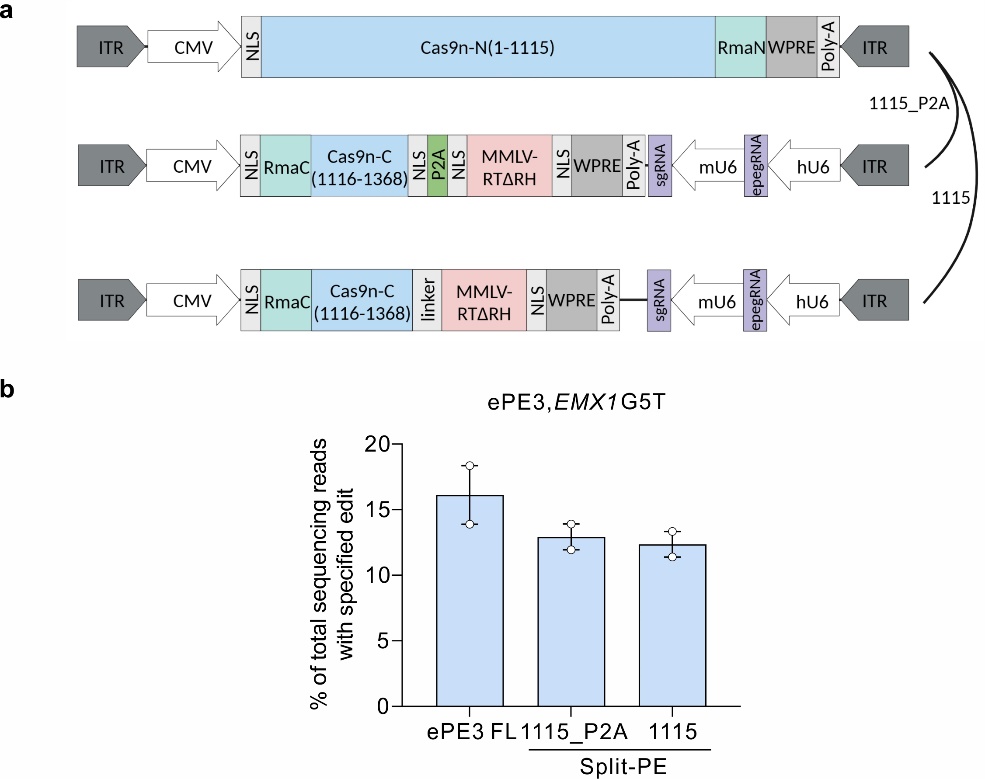

Figure S3. split-PE2-1115s with tethered and untethered MMLV-RTΔR function with similar efficiencies in human HEK293T cells. **a**, Schematic of the AAV vector genomes for split-ePE3-1115 and split-ePE3-1115(P2A). Dissociative MMLV-RTΔR was expressed by substituting the linker between SpCas9n and MMLV-RTΔR with P2A peptide. **b**, The editing efficiencies of split-ePE3-1115 and split-ePE3-1115(P2A) were compared at *EMX1* site. The frequencies of total sequencing reads with specified editing were quantified by NGS and analyzed by CRISSPResso2 (<https://github.com/pinellolab/CRISPResso2>). HEK293T cells were transfected with split-ePE3 or full length ePE3 plasmids at equal molar concentrations. Data are shown as mean ± SEM, n = 2.

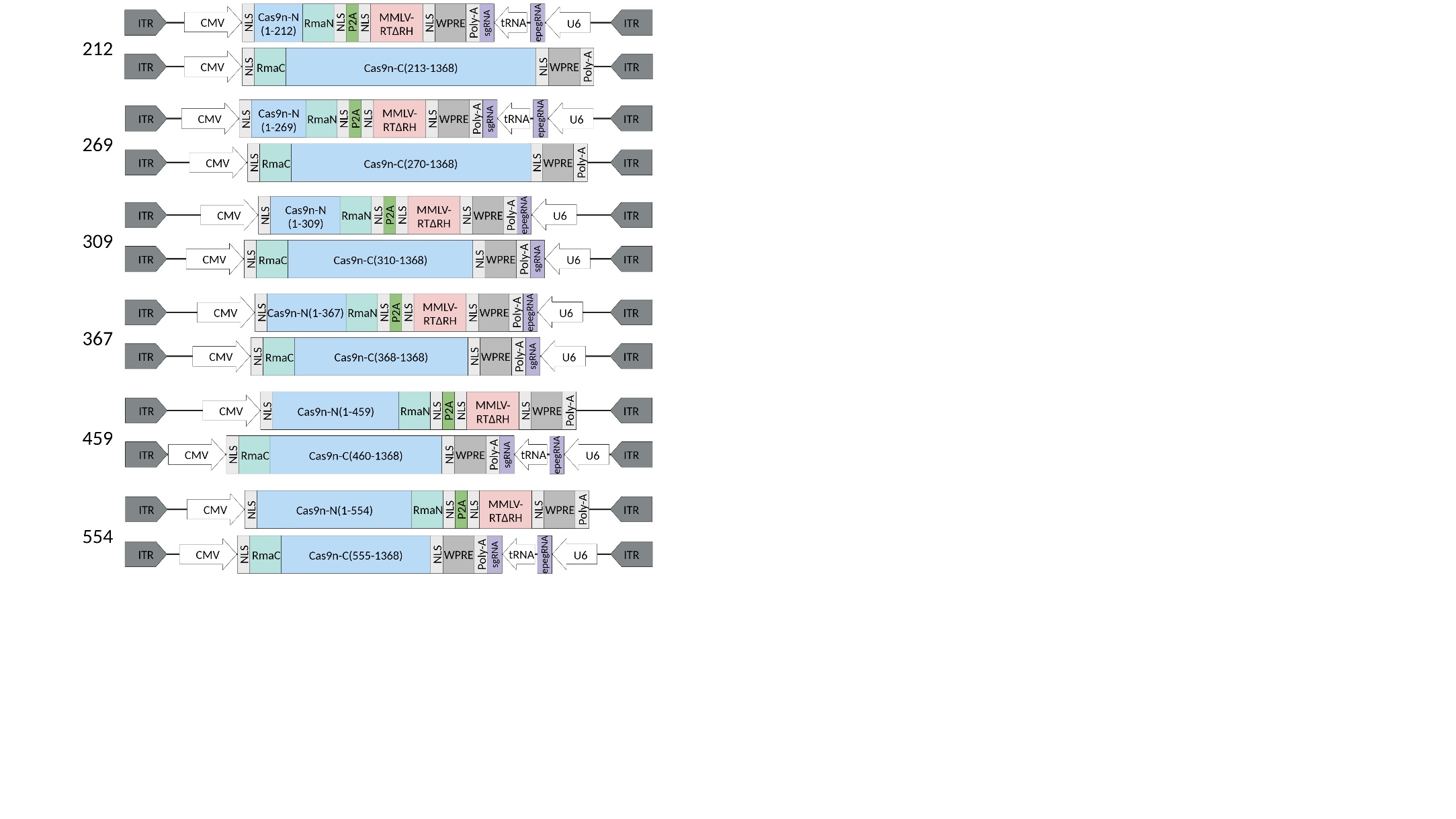

Figure S4. Schematic of the AAV vector genomes for the split-PEs at the first half of SpCas9n. P2A peptide was used to co-express an untethered MMLV-RTΔRH protein.

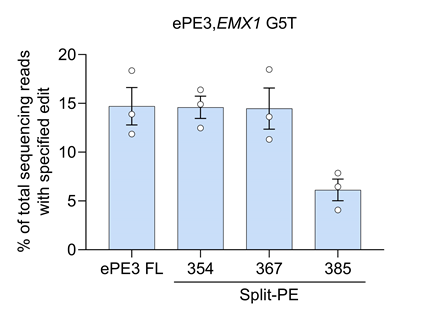

Figure S5. Screening for efficient split sites around split site 367. The frequencies of total sequencing reads with specified editing were quantified by NGS and analyzed by CRISSPResso2 (<https://github.com/pinellolab/CRISPResso2>). HEK293T cells were transfected with split-ePE3 or full length ePE3 plasmids at equal molar concentrations. Data are shown as mean ± SEM, n = 3.

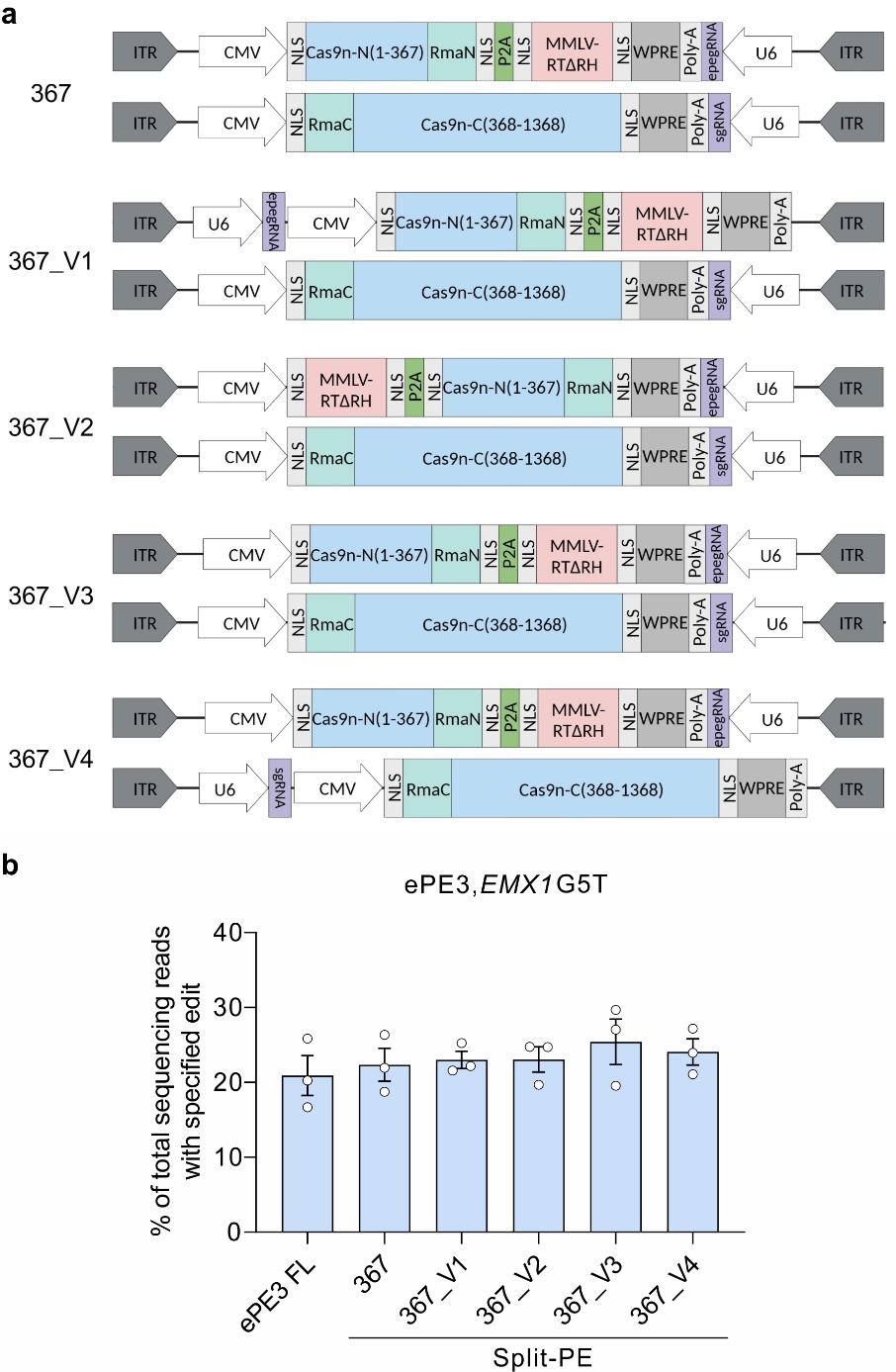

Figure S6. Optimization of split-ePE3-367 by adjusting orientations and positions of pegRNA and nicking sgRNA. **a**, Schematic of different versions of the AAV vector genomes for split-ePE3-367. **b**, Editing efficiencies of different versions of the AAV vector genomes for split-ePE3-367. The frequencies of total sequencing reads with specified editing were quantified by NGS and analyzed by CRISSPResso2 (<https://github.com/pinellolab/CRISPResso2>). HEK293T cells were transfected with split-ePE3 or full length ePE3 plasmids at equal molar concentrations. Data are shown as mean ± SEM, n = 3.

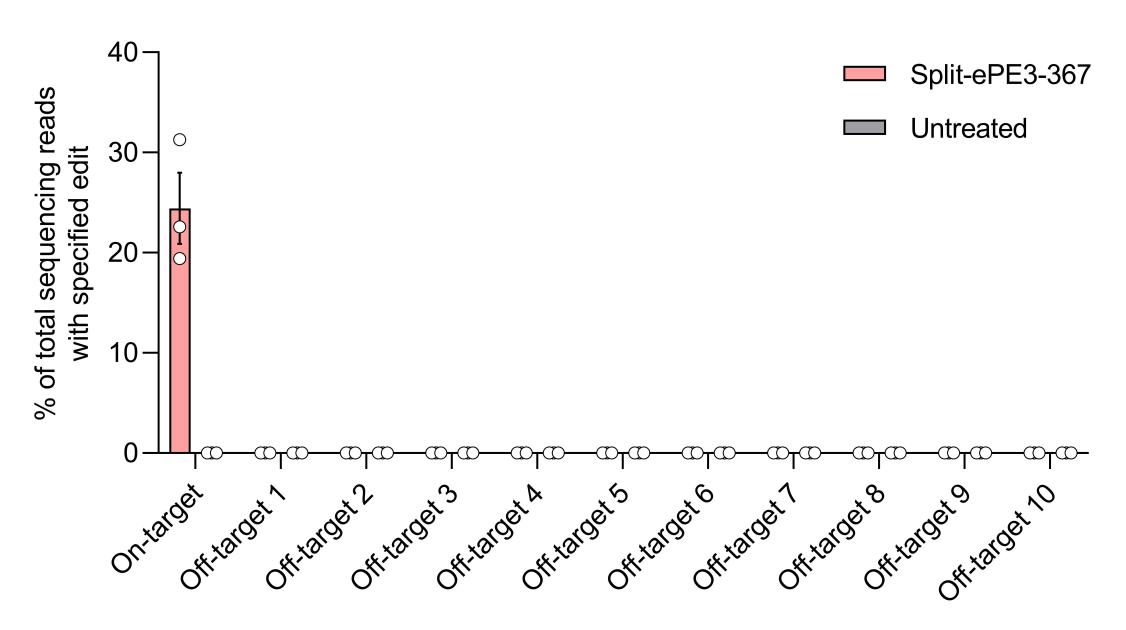

Figure S7. Off-target prime editing for 10 off-target loci (Off-target1–Off-target10) for pegRNA from the liver genomic of split-ePE3-367-AAV-treated and untreated mice. Dots represent individual mice, and error bars represent mean ± SEM of n = 3 mice. Untreated mice were used as negative controls.
